# *Mediocremonas mediterraneus*, a New Member within the Developea

**DOI:** 10.1101/2020.05.12.090621

**Authors:** Bradley A. Weiler, Elisabet L. Sà, Michael E. Sieracki, Ramon Massana, Javier del Campo

## Abstract

The Stramenopiles are a large and diverse group of eukaryotes that possess various lifestyles required to thrive in a broad array of environments. The stramenopiles branch with the alveolates, rhizarians, and telonemids, forming the supergroup TSAR. Here, we present a new genus and species of aquatic nanoflagellated stramenopile: *Mediocremonas mediterraneus*, a free-swimming heterotrophic predator. *M. mediterraneus* cell bodies measure between 2.0-4.0 μm in length and 1.2-3.7 μm in width, possessing two flagella and an oval body morphology. The growth and grazing rate of *M. mediterraneus* in batch cultures ranges from 0.68 to 1.83 d^-1^ and 1.99 to 5.38 bacteria h^-1^, respectively. *M. mediterraneus* was found to be 93.9% phylogenetically similar with *Developayella elegans* and 94.7% with *Develorapax marinus*, two members within the class Developea. The phylogenetic position of the Developea and the ability of *M. mediterraneus* to remain in culture makes it a good candidate for further genomic studies that could help us to better understand phagotropy in marine systems as well as the transition from heterotrophy to phototrophy within the stramenopiles.

## INTRODUCTION

Stramenopiles, also known as Heterokonts (Cavalier-Smith 1986), are a diverse group of eukaryotes found in marine, limnetic and terrestrial systems (Andersen 2004; Massana et al. 2004, 2014; Simon et al. 2015). The stramenopiles branch with Alveolata, Rhizaria, and Telonemia forming a clade supergroup termed TSAR (Adl et al. 2019; Strassert et al. 2019). Within the stramenopiles there are two large monophyletic clades Bigyra and Gyrista, that each contain two monophyletic groups Opalozoa/Sagenista and Ochrophyta/Oomycota, respectively (Derelle et al. 2016). However, these taxonomic classifications are prone to change due to molecular environmental surveys that uncover the expansive diversity previously impossible to characterize by morphological and culturing approaches. These molecular surveys using environmental DNA have uncovered an array of uncultured eukaryotes such as the marine stramenopiles (MAST) clades from diverse aquatic habitats, including also limnetic ecosystems despite their name (Massana et al. 2014).

Stramenopiles have various lifestyles, such as photoautotrophic (most of the ochrophytes, eg. diatoms), heterotrophic predators (e.g. bicosoecids), osmotrophic (e.g. oomycetes) and parasitic (e.g. *Blastocystis*) (Arndt et al. 2000; Andersen 2004; Tan 2008). In some cases, certain stramenopiles have a combination of both photoautotrophic and heterotrophic feeding behaviours, termed mixotrophic that both possess chloroplasts and may also phagocytize prey (Jürgens and Massana 2008). Heterotrophic flagellates capture their prey by raptorial feeding, and prey items are subsequently consumed by phagocytosis (Hansen et al. 1994; Jürgens and Massana 2008). The functional and numerical responses, as discussed in detail by Weisse et al. (2016), are models used to predict the functional ecology of prey consumption by mixo- and heterotrophic aquatic protists. These responses depict how the flagellate growth and grazing rates vary with prey abundance, and can generally be adjusted to Michaelis-Menten kinetics (Weisse et al. 2016).

A novel class within the superphylum Stramenopiles was suggested, termed Developea (Aleoshin et al. 2016), which is composed of biflagellated non-amoeboid heterotrophic nanoflagellates (Cavalier-Smith 2018). Developea was classified under the subphylum Bigyromonada (that includes Pirsonea), within the phylum Gyrista. Within Developea, *Developayella elegans* is a free-swimming aquatic heterotrophic nanoflagellate predator which was previously purported to be grouped within the Oomycetes (AKA Pseudofungi) (Leipe et al. 1996; Cavalier-Smith and Chao 2006). *D. elegans* was recently observed to group with a novel species, *Develorapax marinus*, both forming the Developea class (Aleoshin et al. 2016). Despite not being very abundant in the environment based on molecular surveys, Developea is a very interesting group because of its phylogenetic position. Recent phylogenomic studies have placed *D. elegans* at the base of the Ochrophyta (Leonard et al. 2018), so Developea could represent a key group to understand the transition from heterotrophy to phototrophy within the Stramenopiles. In a previous culturing effort (del Campo et al. 2013), we isolated several heterotrophic nanoflagellates, and one of them remained poorly characterized. Here, we focus on that culture and present a new genus and species of aquatic heterotrophic nanoflagellate: *Mediocremonas mediterraneus*, that clustered closely with both *D. elegans* and *D. marinus* within the Developea class in our 18S rRNA gene phylogenetic analyses. In addition, we present the functional and numerical response of *M. mediterraneus*.

## MATERIAL AND METHODS

### Isolation by single cell sorting

The HNF described here was isolated in a previous culturing effort (del Campo et al. 2013), and the isolation procedure is explained below. Seawater from the Blanes Bay Microbial Observatory sampled on September 30^th^, 2008 was filtered through 3 μm and sent to Bigelow Laboratory for Ocean Sciences (Boothbay Harbor, ME, USA) for cell sorting in a MoFlo™ Flow Cytometer (Dako-Cytomation, Denmark). Digestive vacuoles of heterotrophic protists were stained using the vital stain LysoTracker^®^ (Life Technologies, NY, USA) and cells were sorted based on their green fluorescence (LysoTracker^®^ fluorescence) and the absence of chlorophyll fluorescence. Details of the staining protocol and flow cytometer setup are described elsewhere (Rose et al. 2004; Heywood et al. 2011). Side scatter was used to select only the smallest protists, approximately <10 μm in diameter. Individual target cells were deposited into 24-well plates, in which some wells were dedicated for positive controls (10 cells per well) and negative controls (0 cells per well). All wells on the microplates contained 1ml aged seawater together with natural bacteria at a final concentration of 5 × 10^6^ bacteria ml^-1^. Multi-well plates were hand carried back to the Institut de Ciències del Mar (Barcelona, Catalonia, Spain) by plane at room temperature (12 hours).

### Culture maintenance

Cultures were maintained in 50 mL flasks and transferred every two to four weeks to fresh media (aged sea water with added bacteria) at 1/10 dilution. The full culturing protocol is outlined in del Campo *et al*. (2013), however in short batch cultures were prepared by adding small aliquotes of the *M. mediterraneus* culture to sterile seawater (about 2 liters) containing *Dokdonia donghaensis* MED134 at varying cell abundances. Bacterial and flagellate counting was conducted once or twice a day using epifluorescence microscopy and DAPI stain by fixing 3 mL aliquots of the batch culture with glutaraldehyde directly after sampling. Growth rate was calculated from the exponential increase of flagellate cells. Grazing rate was calculated based on the calculations by Frost (1972) using the resultant growth rates, the exponential decrease of bacterial cells and the geometric average concentration of flagellates and bacteria during the incubation (Frost 1972). Growth efficiency was calculated from the ratio of protist biomass produced as compared with bacterial biomass consumed.

### Sequencing and phylogenetic analysis

For molecular analysis, the whole culture was filtered on 0.6 μm polycarbonate filters, DNA was extracted by standard procedures, and 18S rDNA genes were PCR amplified with eukaryote-specific primers (Díez et al. 2001). Complete sequences of 18S rDNA were obtained with five internal primers by the MACROGEN Genomics Sequencing Services (accession numbers JX272636 & XXXXXXXX). The longest sequence of *M. mediterraneus* (JX272636), a subset of previously published Stramenopiles and two outgroups were aligned using MAFFT v7.453 (setting: -auto) (Katoh and Standley 2013). Regions of poor alignment were removed using trimAl v1.4 (Capella-Gutierrez et al. 2009) with a gap and similarity threshold of 0.3 and 0.001, respectively (settings: -gt 0.3 -st 0.001). Finally, to explore *M. mediterraneus* phylogeny, maximum likelihood (ML) inference was conducted with RAxML v8.2.12 (Stamatakis 2014) assuming the GTRCAT model using rapid bootstrap analysis for 1,000 iterations.

### Scanning Electron Microscopy

Scanning Electron Microscopy was performed as in Garcés et al. (2006). Samples were fixed for 2 h in 2% OsO_4_ diluted in seawater, or with 2% glutaraldehyde. Cells were subsequently washed with distilled water (2 h) and filtered onto a 0.6 μm pore size Nuclepore™ filter (Whatman, Maidstone, UK). Samples were dehydrated in an ethanol series (30, 50, 70, 96, and 100%) for 15 min each, followed by an acetone series (25, 50, 75, and 100%) for 15 min each. Samples were critical point-dried in liquid CO_2_ using a BAL-TEC CPD 030 critical point drying apparatus (Balzers Union, Balzers, Germany). Filters were subsequently glued to SEM-stubs with colloidal silver, sputter coated with gold palladium, and examined with a Hitachi S-3500N (Nissei Sangyo Co. Ltd., Tokyo, Japan) SEM operating at 5 kV.

## RESULTS AND DISCUSSION

In terms of gross morphology *M. mediterraneus* is relatively similar to the other members of the Developea clade (Aleoshin et al. 2016; Tong 1995). Ovoid cell bodies measure 2.0 – 4.0 μm in length and 1.2 – 3.7 μm in width (Fig. 1A) and possesses a depression on the anterior ventral surface (Fig. 1B, 1D). Protruding from the depression are two flagella of equal length, unlike those of both *D. elegans* and *D. marinus*, measuring roughly twice the cell body length. The anterior flagellum possesses mastigonemes and projects from the cell in an arc (Fig. 1). The posterior flagellum projects posteriorly out from the cell depression relatively straight, not possessing mastigonemes. The function of the Developea flagella have been previously described for both locomotion (posterior) and current generation (anterior) to aid in prey capture (Aleoshin et al. 2016; Tong 1995).

**Figure 1:**
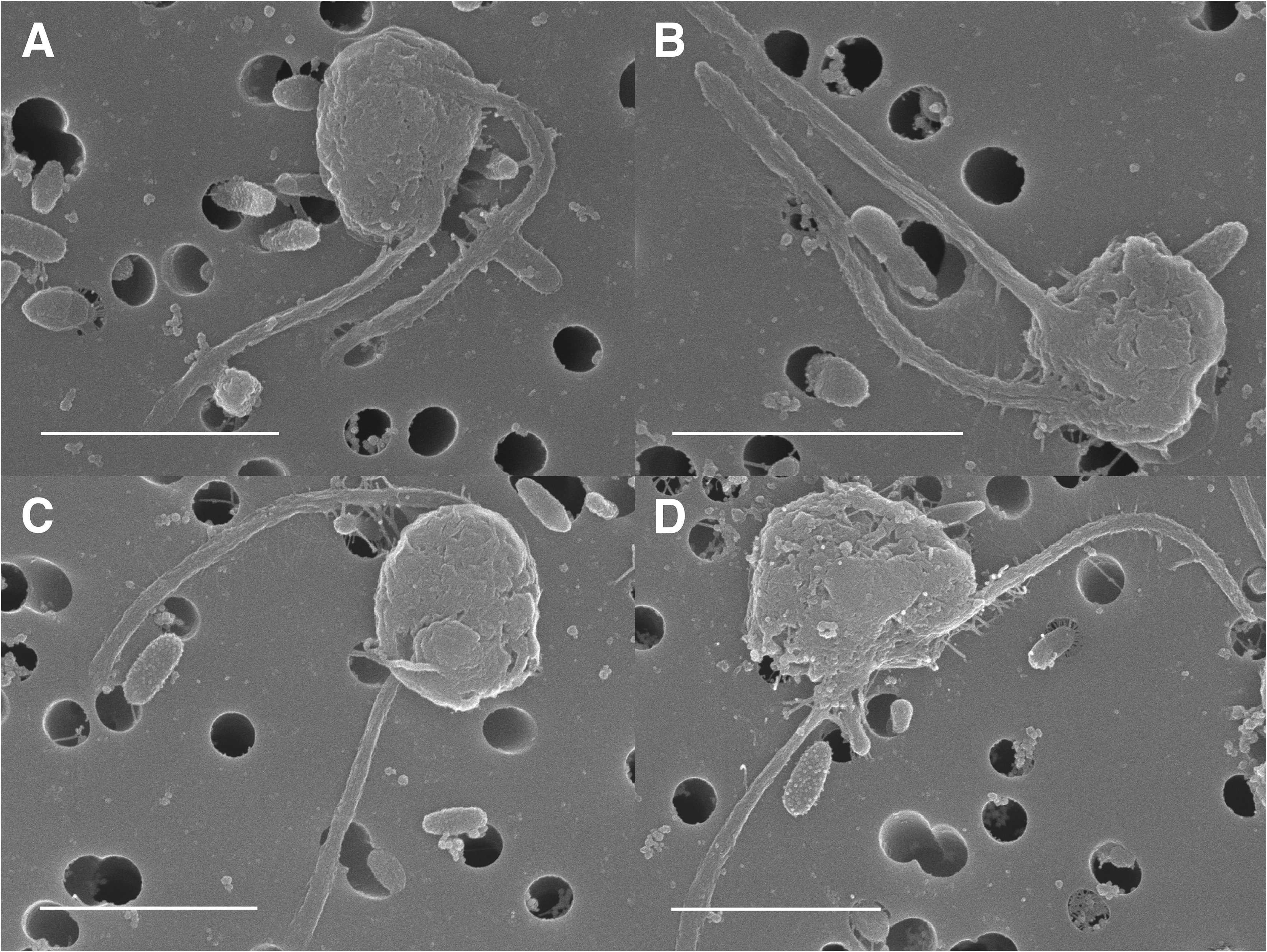
SEM images of *M. mediterraneus* cells at varying angles (**A-D**), showing an ovoid body morphology and two flagella protruding from a surface depression ventrally located on the anterior portion of the cell. Anterior flagellum typically held in an arc possessing tubular mastigonemes. Scale bars measure 3 μm.

Sequences were analyzed using BLAST for phylogenetic similarity at the 18S rRNA gene; it was 93.9% similar with *Developayella elegans* (Leipe et al. 1996) and 94.7% with *Develorapax marinus* (Aleoshin et al. 2016). When analyzing the 18S rRNA gene sequence of *M. mediterraneus* using maximum likelihood (ML), there is supported clustering with previously described members of the Developea class (Leipe et al. 1996; Aleoshin et al. 2016), as well as supported clustering with two other uncultured eukaryotes within the class Developea, both retrieved from deep-sea sediments (Fig. 2). The next closest cluster observed in the ML tree is the “Abyssal” group (as in Aleoshin et al. 2016) including eukaryotes found in the Pacific abyssal plains (Takishita et al. 2010). Additionally, sister clade to the Developea+Abyssal grouping are the Pseudofungi from the phylum Gyrista. As mentioned in the introduction the most recent phylogenomic analysis of this part of the eukaryotic tree of life shows *D. elegans* at the base of the Ochrophyta (Leonard et al. 2018), however this position is not strongly supported in our 18S rDNA tree and in other phylogenetic reconstructions (Aleoshin et al. 2016). Obtaining in the future the genome of *M. mediterraneus* can help to resolve this area of the Stramenopiles tree within Ochrophyta and Pseudofungi.

**Figure 2:**
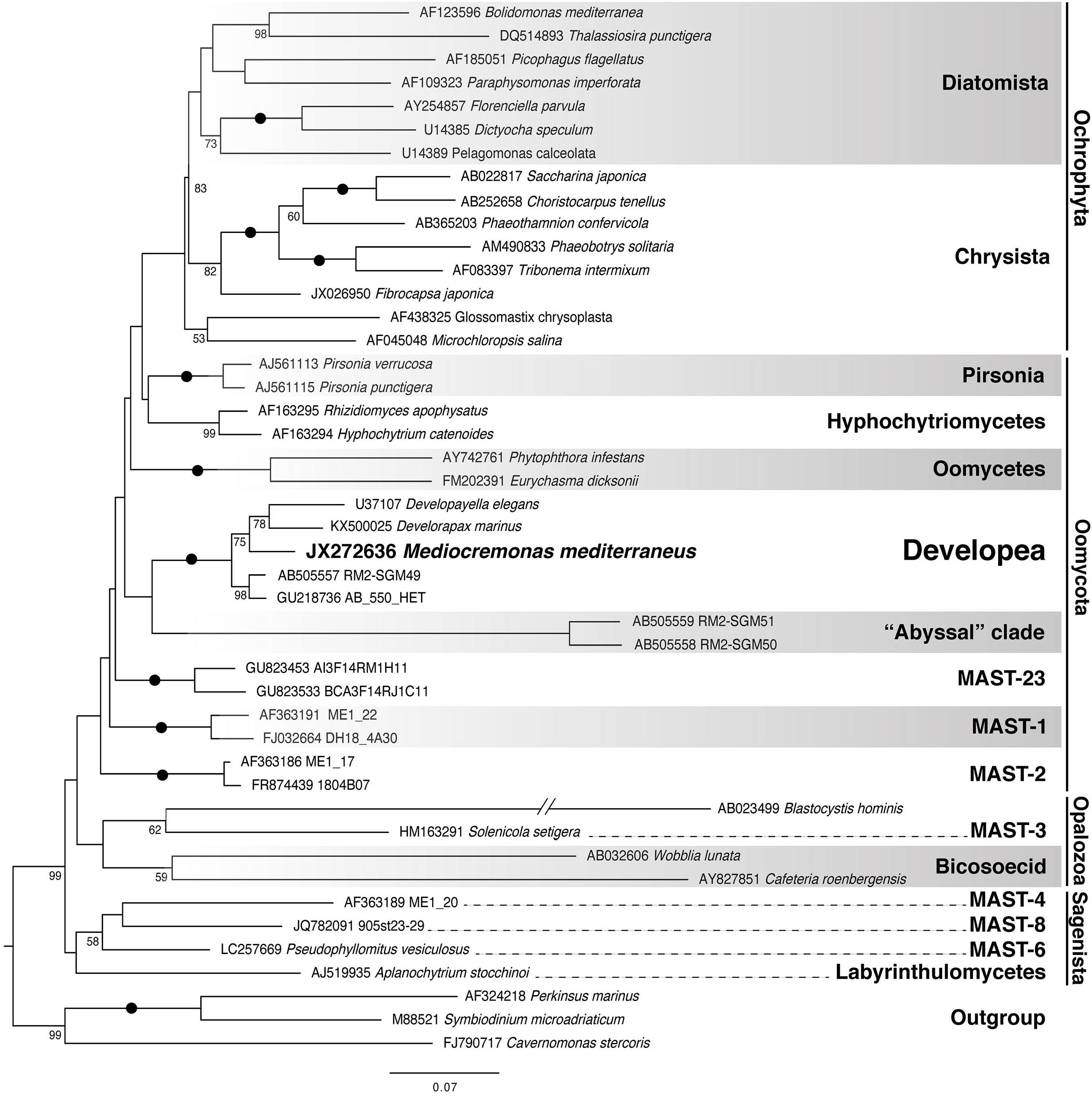
Maximum likelihood phylogenetic tree with partial (~1700 bp) 18S rRNA gene sequences showing the clustering of *M. mediterraneus* with the Developea group within the Gyrista clade. ML tree based off 1,000 iterations and only bootstrap values > 50 are shown. The symbol (•) represents bootstrap values of 100.

We prepared several batch cultures where we added an aliquote of *M. mediterraneus* on flasks containing varying cell abundances of bacterial strain and followed the dynamics over time of both components (Fig. S1). By following the exponential predator increase and prey decrease, we calculated the growth rate and the grazing rate in each of these batch cultures. The grazing rate of*M. mediterraneus* ranged from 1.99 to 5.38 bacteria h^-1^, and the growth rate ranged from 0.68 to 1.83 d^-1^ (Table 1). The yield of bacteria to flagellate (Yield_b/f_) ranged between 54.8 and 84.1 (mean of 74.8, SD = 11.3). The doubling time in days observed for *M. mediterraneus*, ranging from 0.38 to 1.02, was relatively longer than that of other heterotrophic nanoflagellates (HNF), as *Paraphysomonas imperforata, Minorisa minuta*, and *Cafeteria roenbergensis* have doubling times of 0.28, 0.44, and 0.45, respectively (Goldman and Caron 1985; del Campo et al. 2013; De Corte et al. 2019). The functional and numerical responses showed a relatively clear trend (Fig. 3), wherein the growth and grazing rates increase as bacterial prey abundance increases, up to a theoretical asymptotic maximum in terms of grazing rate (I_max_) and growth rate (R_max_), inferred to be 6.23 (h^-1^) and 2.42 (d^-1^), respectively (Fig. 3).

**Figure 3:**
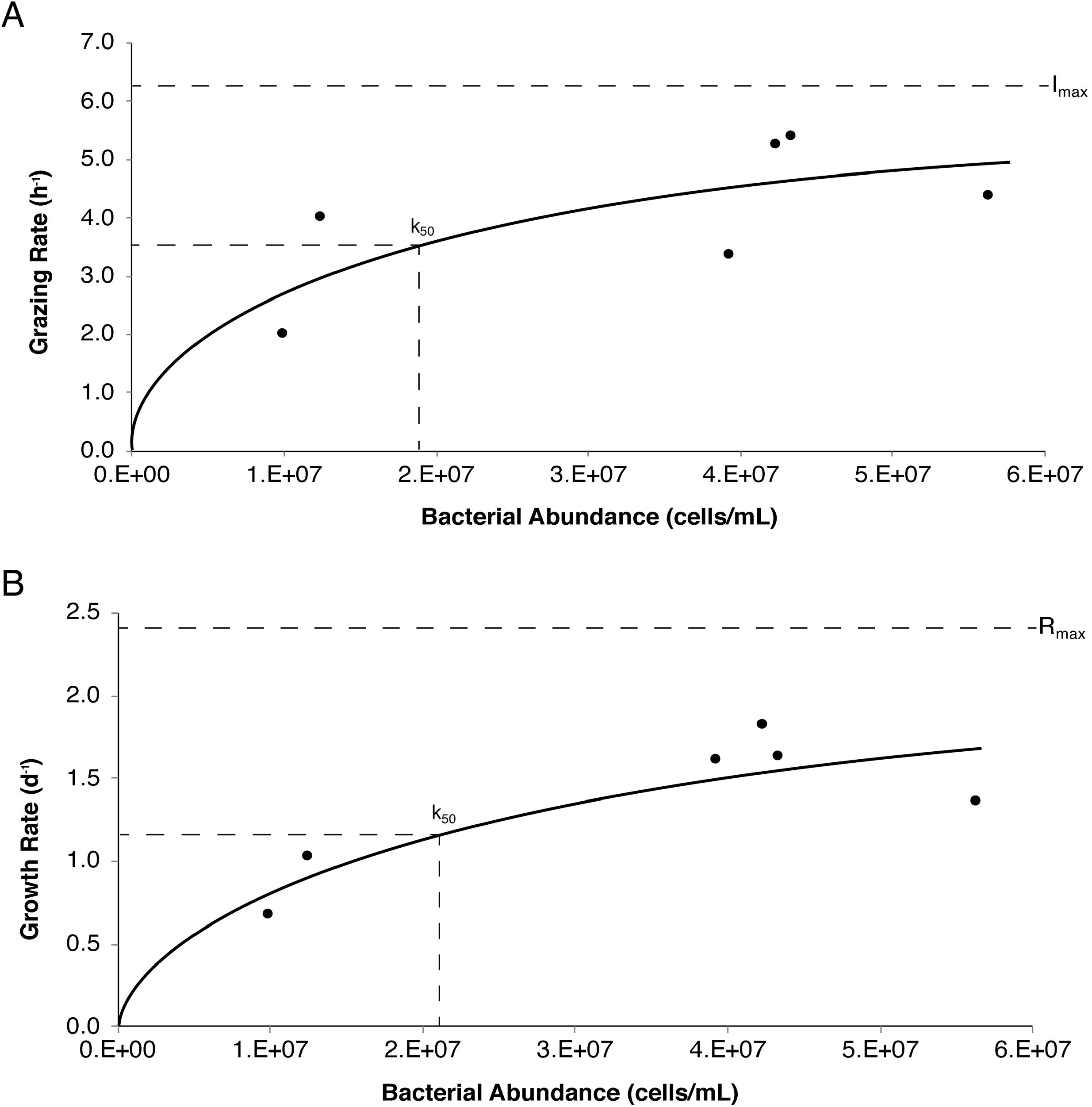
Numerical and functional responses of *M. mediterraneus*. **A.** showing the specific grazing rate (h^-1^) on the y-axis as a response to increased bacterial abundance (cells/mL), with I_max_ indicating the theoretical maximum grazing rate and the resultant k_50_. **B.** showing the specific growth rate (d^-1^) on the y-axis, as a response to increased bacterial abundance (cells/mL), with R_max_ indicating the theoretical maximum growth rate and resultant k_50_.

**Table 1:** Growth and bacterial grazing characteristics of *M. mediterraneus* by samples (1-4). Samples 1 & 2 have replicates denoted by A & B.

| Sample | Date | Initial<br>Bacterial<br>Conc.<br>(cell/mL) | Growth<br>Rate<br>(d<sup>-1</sup>) | Doubling<br>Time<br>(days) | Grazing<br>Rate<br>(h<sup>-1</sup>) | Yield<br>(bac/fla) | Growth<br>Efficiency<br>(Yield)<br>(%) |
| --- | --- | --- | --- | --- | --- | --- | --- |
| 1A | Oct. 2009 | 9.86E+06 | 0.68 | 1.02 | 1.99 | 70.90 | 35.31 |
| 1B | Oct. 2010 | 5.62E+07 | 1.37 | 0.51 | 4.38 | 82.94 | 30.19 |
| 2A | Nov. 2009 | 4.33E+07 | 1.64 | 0.42 | 5.38 | 83.04 | 30.15 |
| 2B* | Nov. 2009 | 3.92E+07 | 1.62 | 0.43 | 3.37 | 54.81 | 45.68 |
| 3 | May 2011 | 1.24E+07 | 1.04 | 0.67 | 4.01 | 72.78 | 34.40 |
| 4* | Dec. 2011 | 4.22E+07 | 1.83 | 0.38 | 5.24 | 84.06 | 29.79 |

\* with natural viruses

Bacterivory in aquatic ecosystems is a crucial functional role played predominantly by pico- and nanoflagellates (up to 5 μm) to maintain bacterial populations (Sherr and Sherr 2002). Grazing on bacteria is necessary for nutrient recycling by releasing waste products back into the environment in the form of inorganic compounds (such as iron, ammonium and phosphate), and particulate and dissolved organic compounds (Sherr and Sherr 2002; Massana et al. 2009). The grazing rate of *M. mediterraneus* ~4.1 bacteria h^-1^ is greater than some HNFs such as the closely related MAST-1C group (Fig. 2), which has a grazing rate of 3.6 bacteria h^-1^ and the more distantly related MAST-4 group, 1.5 bacteria h^-1^ (Table 2). In contrast, the grazing rate of *M. mediterraneus* is much lower than several other cultured HNFs, including various gluttonous members of the genus *Bodo spp*. (34.3 – 160 bacteria h^-1^), and members of the genera *Spumella sp*. and *Ochromonas sp*. (37 and 63 bacteria h^-1^, respectively) (Table 2). In general, grazing rates of cultured HNFs (Eccleston-Parry and Leadbeater 1994), which are suggested to be poor models of natural and dominant taxa (Massana et al. 2009), are much higher than community grazing rates typically between 2 and 20 bacteria h^-1^ (Jürgens and Massana 2008).

**Table 2:**
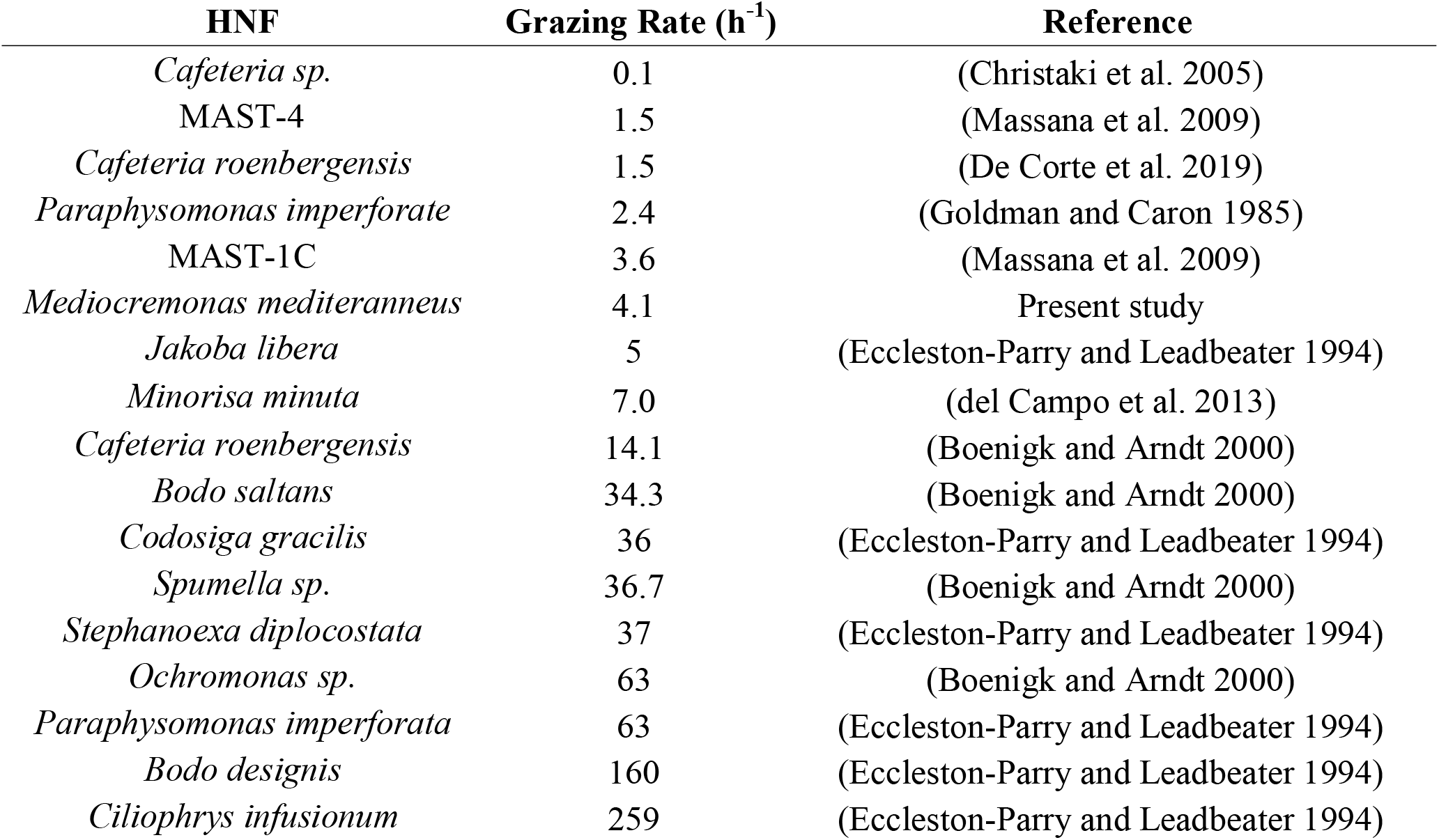
Comparison of heterotrophic nanoflagellate (HNF) grazing rates ranked lowest to highest.

Blanes Bay is an oligotrophic coastal area of the NW Mediterranean and resides within proximity to a continental submarine canyon. The microbial food web dynamics of Blanes Bay, similar to other regions, are largely influenced by environmental factors (i.e. salinity, Chl a, temperature, nutrients, etc…) and may change given a shift in just one of those factors (e.g. increased sea surface temperature, Vázquez-Domínguez et al. (2012)). Given the oligotrophic nature of Blanes Bay, and considering the low abundance of *M. mediterraneus* (Giner et al. 2019), it may be present perhaps as opportunistic consumers in microbial “hotspots” that arise in the oligotrophic ecosystem (Stocker et al. 2008; Dang and Lovell 2016).

The ability to have *M. mediterraneus* in culture and its interesting phylogenetic position make this organism a good candidate for further genomic studies that could help us to better understand phagotropy in marine systems as well as the transition from heterotrophy to phototrophy within the stramenopiles.

## TAXONOMIC SUMMARY

**Phylum Gyrista, Cavalier-Smith, 1997**

**Class Developea, Aleoshin, 2016**

**Order Developayellales, Cavalier-Smith, 1997**

**Family Developayellaceae, Cavalier-Smith, 1997**

**Genus *Mediocremonas, del Campo & Weiler, 2020***

**Species *Mediocremonas mediterraneus, del Campo & Weiler, 2020***

### *Mediocremonas* n. gen. del Campo & Weiler

**Zoobank ID:** XXXXX

**Diagnosis:** Heterotrophic oval-shaped protist possessing two flagella with a free-swimming predatorial feeding behaviour. Similar to other genera in the Developea (Aleoshin et al. 2016; Tong 1995), *Mediocremonas* flagella are ventrally located, with the anterior flagellum held in an arc and the posterior flagellum projecting posterior from a conspicuous depression. The anterior flagellum is observed to possess mastigonemes while generating current by performing a slow sweeping motion and the posterior beats rapidly for locomotion. The genus type species, *M. mediterraneus*, was isolated from Blanes Bay (Catalonia, Spain) and the 18S sequence was observed to be 93.9% similar to *Developayella elegans* from surface estuarine water of Southampton Water, UK (Leipe et al. 1996) and 94.7% similar to *Develorapax marinus* from coastal waters of the Red Sea (Aleoshin et al. 2016).

**Type species:** *M. mediterraneus* n. sp. del Campo & Weiler

**Etymology:** Mediocre – [average (l)] + monas – [unicellular organism (l)]

### *M. mediterraneus* n. sp. del Campo & Weiler

**Zoobank ID:** XXXXX

**Diagnosis:** Cell bodies measure between 2.0-4.0 μm in length and 1.2-3.7 μm in width, possessing an oval body morphology. The general body size is comparatively smaller than other members of Developea, wherein *Developayella elegans* is roughly twice as large (3.5-8.5 μm in length and 2.0-6.0 μm in width) and the body size of *Develorapax marinus* is relatively largest (7.0-10.0 μm in length and 4.0-6.0 μm in width). Each cell contains two flagella originating from a ventral anterior depression in the cell body. Within the cell depression, the two flagella lay either anterior or posterior relative to one another, both with specific form and function. The anterior flagellum of *Developea* has been previously described to be held in an arc containing mastigonemes that sweeps slowly ventral posterior to the cell, while the posterior beats rapidly for locomotion. The anterior and posterior flagella of *M. mediterraneus* measure between 5.5-6.0 μm in length and 0.2-0.3 μm in width dissimilar to the unequal flagella measurements in both *D. elegans* and *D. marinus*.

**Type material:** Fig. 1 from cultures 9 and 11, isolated from Blanes Bay Microbial Observatory (NW Mediterranean Sea) and maintained at the Institut de Ciències del Mar (Barcelona, Catalonia, Spain). Scanning microscopy samples are also preserved there.

**Type sequence:** The SSU rRNA gene sequence is Genbank JX272636

**Type locality:** Blanes Bay (Catalonia, Spain)

**Etymology:** mediterraneus [from the Mediterranean (l)]

## Supporting information

Supplementary Figure S1: Cell abundance dynamics of M. mediterraneus (black) and bacterial prey (grey) in batch culture experiments (1 to 4). Experime

## ACKNOWLEDGEMENTS

This work was supported by two grants from the Spanish government, FLAME (CGL2010-16304, MICINN) and ALLFLAGS (CTM2016-75083-R, MINECO), and by the National Science Foundation award DEB-1031049 to MES. BW was supported by the Natural Sciences and Engineering Research Council of Canada Postgraduate Scholarships-Doctoral program (NSERC PGS-D). BW and JdC were supported by startup funds from University of Miami, Rosenstiel School of Marine and Atmospheric Sciences.

---

**Supplementary Figure S1:** Cell abundance dynamics of *M. mediterraneus* (black) and bacterial prey (grey) in batch culture experiments (1 to 4). Experiments 1 & 2 have two replicates indicated by A & B.

## Notes

### Competing Interest Statement

The authors have declared no competing interest.

### Summary of Updates

Author information revised and updated

