## Supplementary figures and images for "*Mediocremonas mediterraneus*, a New Member within the Developea"

### Supplementary Figure S1: Cell abundance dynamics of M. mediterraneus (black) and bacterial prey (grey) in batch culture experiments (1 to 4). Experime

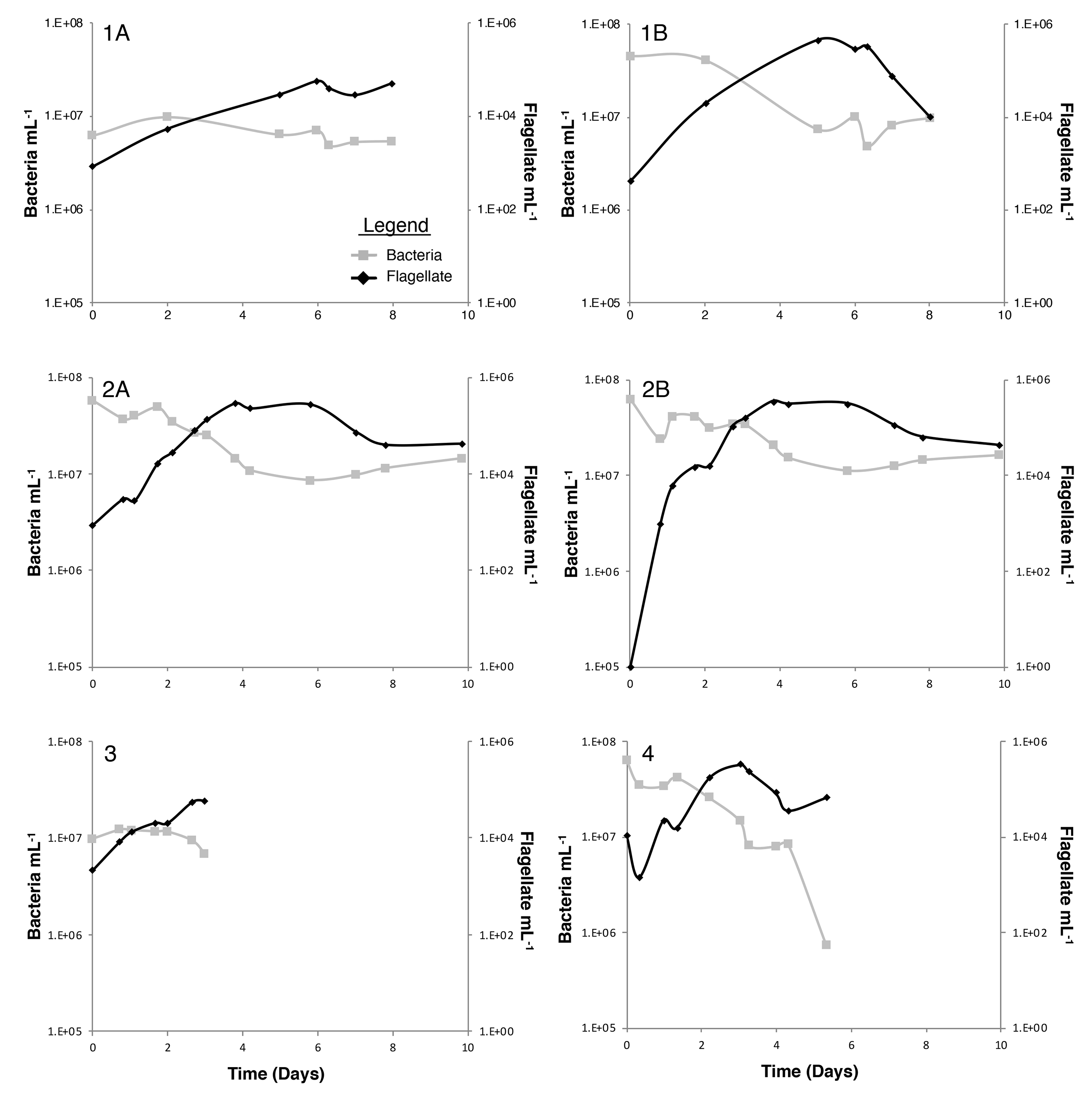
